# Latitudinal gradients in stable hydrogen isotopes of North American trees

**DOI:** 10.1101/2025.01.29.635453

**Authors:** Frank Keppler, Lars Berger, Markus Greule, Jan Esper, Anna Wieland

## Abstract

Stable hydrogen isotopes (*δ*^2^H values) of plant matter provide information about ambient climate conditions and physiological processes. *δ*^2^H values of tree stem wood methoxy groups (*δ*^2^H_TM_) are increasingly used to reconstruct the isotope composition of precipitation (*δ*^2^H_precip_) and temperature variability in mid latitudes regions. However, empirical evidence of the basic latitudinal gradients in North America and their inter-continental context are still missing. We sampled 104 tree cores in 13 North American sites ranging from 38°-69°N and investigated the relationship between *δ*^2^H_TM_ and *δ*^2^H_precip_ values. Recorded *δ*^2^H_TM_ values are highly depleted in ^2^H relative to meteoric water and align closely with predictions from a biochemical model. Reconstructed temperature changes, derived from *δ*^2^H_TM_ values, agree well with instrumental data in North American since 1932. Our results reveal that the latitudinal gradients of *δ*^2^H_TM_ values closely resemble *δ*^2^H_precip_ values across North America. When combining these spatial patterns with *δ*^2^H_TM_ and *δ*^2^H_precip_ data from Europe a strong correlation (R^2^ = 0.94, p < 0.001) and a consistent fractionation of –193 ± 13 mUr, representative for Northern Hemisphere mid-to-high latitude sites, is established. The validation across North American and European transects serves as a foundational basis for advancing applications in biogeochemistry and paleoclimate research.

## Introduction

The distinct pattern of stable hydrogen isotope (*δ*^2^H) values of plant methoxy groups (R-OCH_3_) is gaining recognition as a proxy across diverse research fields. Recent studies have utilized the isotopic signatures of plant methoxy groups for various applications, including tracing the geographical origin of potato tubers (Keppler & Hamilton, 2008), authenticating vanillin (Greule *et al*., 2010; Geißler *et al*., 2017; Wilde *et al*., 2019), assessing water loss in fruits and vegetables during storage (Greule *et al*., 2015), determining the origin of plant-related methane emissions (Keppler *et al*., 2008) and chloromethane emissions (Keppler *et al*., 2004; Derendorp *et al*., 2011; Keppler *et al*., 2020; Hartmann *et al*., 2023), characterizing lignin sources, and investigating the origin of coal-bed methane (Keppler, 2021; Lloyd *et al*., 2021; Cox *et al*., 2024a). Due to the temporal stability of OCH_3_ and the distinct isotopic signatures among different plant tissues, this isotopic tool recently been applied to aid in historical apportionments of particulate organic matter and sediment sources (Cox *et al*., 2024a; Cox *et al*., 2024b). Plant methoxy isotopic patterns have been used as a proxy for photorespiration processes in trees (Lloyd *et al*., 2023) and to demonstrate that the serine phosphate pathway is part of a ’photosynthetic C_1_ pathway’ (Jardine *et al*., 2024).

*δ*^2^H values of tree methoxy groups (*δ*^2^H_TM_) were also applied to record the hydrogen isotopic composition of precipitation (*δ*^2^H_precip_) which is primarily influenced by temperature. As a result, several previous studies have utilized *δ*^2^H values of tree methoxy groups (*δ*^2^H_TM_) as a climate proxy in mid-to-high latitudes. Studies include alpine regions (Gori *et al*., 2013; Riechelmann *et al*., 2017), permafrost forests in northeastern China (Lu *et al*., 2020; Wang *et al*., 2020), Mt. Kilimanjaro (Hepp *et al*., 2017), mid-elevation sites in southern Germany, (Anhäuser *et al*., 2020; Wieland *et al*., 2022), Mediterranean region (Mt. Smolikas, Greece; (Wieland *et al*., 2024), mummified wood specimens from the earliest Eocene (∼55 Ma) kimberlite pipes in the subarctic Northwest Territories of Canada (Anhäuser *et al*., 2018), and Miocene wood (Porter *et al*., 2022).

The strong relationship between *δ*^2^H_precip_ and *δ*^2^H_TM_ was first discovered by Keppler and colleagues to determine the geographical origin of tree samples, particularly in Europe (Keppler *et al*., 2007). The initial isotope fractionation between *δ*^2^H_precip_ and *δ*^2^H_TM_ values – commonly expressed as the apparent isotope fractionation (*ε*_app_) – was originally estimated to be –216 ± 19 mUr, then later revised to –213 ± 17 mUr (Anhäuser *et al*., 2017).

The analytical procedure for *δ*^2^H_TM_ measurements is based on the ‘Zeisel-method’ where the methyl moiety (-CH_3_) of methoxy groups is quantitatively converted into gaseous iodomethane (CH_3_I) (Zeisel, 1885) which is then subsequently measured by gas chromatography high temperature conversion isotope ratio mass spectrometry (GC-HTC-IRMS) (Greule *et al*., 2008). It is important to note, that reliable stable isotope analysis requires appropriate reference materials (RMs) to normalize raw measurement data to internationally comparable isotope scales using primary reference materials (Coplen, 2011; Coleman & Meier-Augenstein, 2014). Importantly, in recent years considerable analytical progress has been made in providing RMs for thoroughly normalizing δ^2^H_TM_ values to the VSMOW scale over a *δ*^2^H range of –100 to – 350 mUr (Greule *et al*., 2019; Greule *et al*., 2020). This advancement now enables improved comparability of *δ*^2^H values in plant methoxy groups across different laboratories. Building on this major achievement the *δ*^2^H_TM_ values of the tree samples from Anhäuser *et al*. (2017) were remeasured and normalized using the newly available RMs (Greule *et al*., 2020; Greule *et al*., 2021), resulting in a slightly revised mean hydrogen isotope fractionation between tree methoxy groups and local precipitation (*ε*_TM/precip_) of −200 ± 14 mUr. In mid-latitudes, where temperature is the primary driver of stable hydrogen isotope signatures in precipitation, this robust correlation between *δ*^2^H_precip_ and *δ*^2^H_TM_ values can be utilized to reconstruct past temperature variations. Based on this relationship and further theoretical biochemical considerations, Greule and colleagues (Greule *et al*., 2021) suggested the following equation to reconstruct mean annual temperature (MAT) from *δ*^2^H_TM_ measurements:

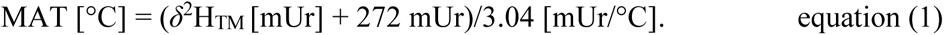

A limitation of the previous dataset is its focus on tree samples along a European transect, making it uncertain whether the observed gradient is applicable to other continents or extendable to broader intercontinental relationships across the Northern Hemisphere. In this paper, we investigate the relationship between *δ*^2^H values of tree methoxy groups and of local precipitation from various geographical locations along a North American transect. We focus on spatial changes from 38-69° N over the recent 30 years. The relative *δ*^2^H_TM_ variations are compared with observed temperature changes across North America considering a range of 90 years (1932–2021) separated in 30-year time intervals. We finally integrate the relationship between *δ*^2^H_TM_ and *δ*^2^H_precip_ values in North America with those from Europe to establish an inter-continental gradient valid over a range ≥ 200 mUr of *δ*^2^H_precip_ values for the Northern Hemisphere and compare these with patterns of *δ*^2^H values in tree cellulose and leaf n-alkanes.

## Materials and methods

### Field sites and tree ring samples

In 2022 and 2023, cores of 104 trees were collected in 13 sampling sites across North America (Fig. 1). Full details of the field sites, tree species and numbers are provided in Table S1. Stem cores were obtained using a 5-mm diameter borer at breast height ∼1.2 to 1.4 m above ground. Tree samples were processed according to standard dendrochronological techniques (Stokes, 1996) and tree ring width (TRW) measured using a high-precision Lintab devise (Rinntech GmbH, Heidelberg, Germany). Stem cores were split into 30 years section from 2021 to 1932 and each section was subsequently homogenized and milled using a ball-mill (Retsch GmbH, Haan, Germany). To establish the intercontinental relationship, we used *δ*^2^H_TM_ values reported by Greule *et al*. (2021) considering data from sites with n > 1.

**Figure 1.**
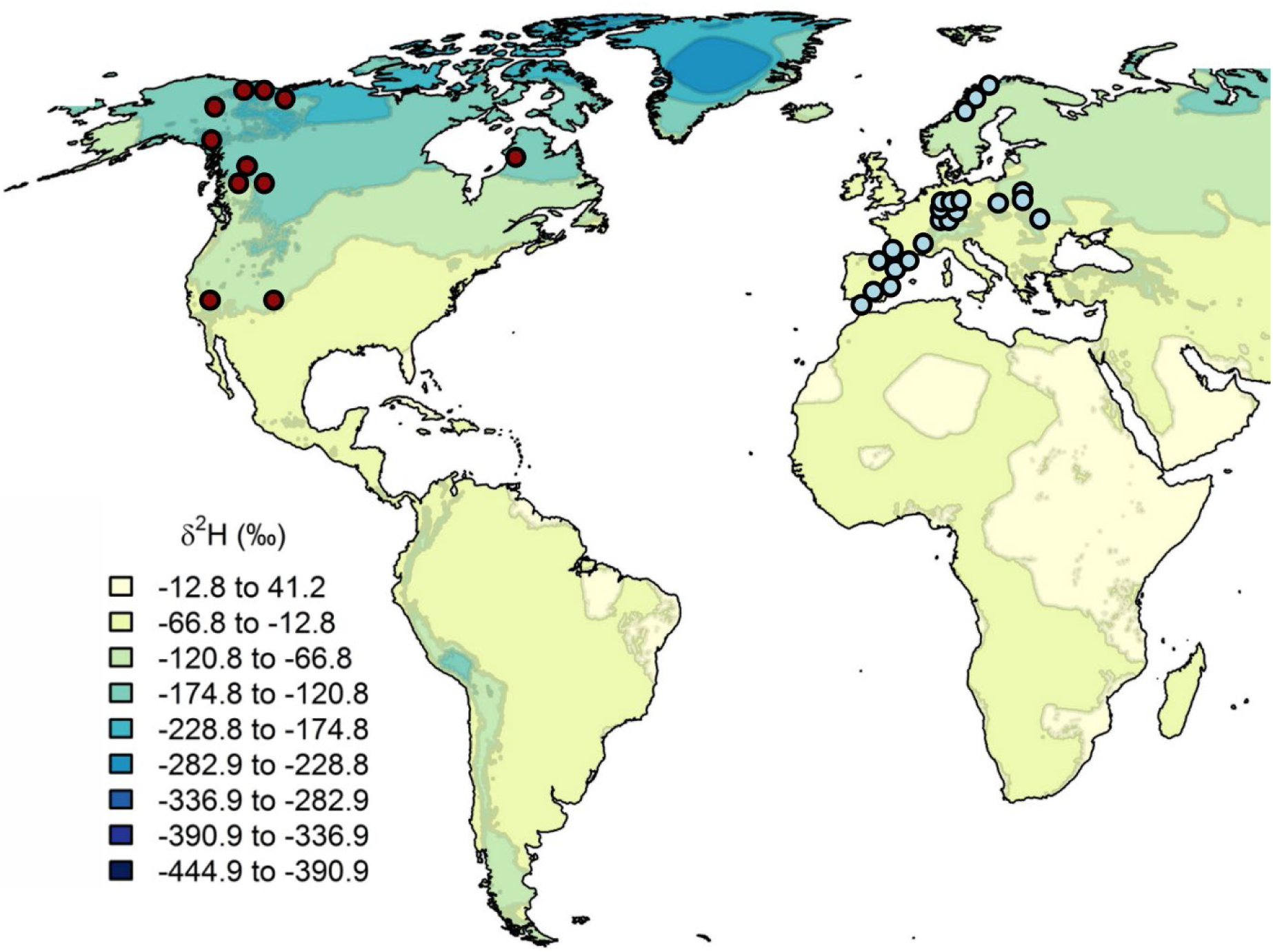
Tree sampling sites along North American (red, this study) and the European transects (blue, from Greule *et al*. (2021)). Contour colors are *δ*^2^H_precip_ values derived from the Online Isotope Precipitation Calculator.

### Stable hydrogen isotope measurements of tree methoxy groups

The *δ*^2^H values of CH_3_I released from the methoxy groups of wood samples and reference materials treated with hydriodic acid (HI) (for detailed description please refer to Greule et al. 2008) were analyzed using GC-HTC-IRMS. The analytical system consists of an A200S auto sampler (CTC Analytics, Zwingen, Switzerland), a HP 6890N gas chromatograph (Agilent, Santa Clara, USA) and a Delta^PLUS^XL isotope ratio mass spectrometer (Thermo Fisher Scientific, Bremen, Germany). The thermo conversion reactor [empty ceramic tube (Al_2_O_3_), length 320 mm, 0.5 mm i.d.] at a reactor temperature of 1450 °C was coupled to the IRMS via a GC Combustion III Interface (ThermoQuest Finnigan, Bremen, Germany). The Zebron ZB-5MS capillary column (Phenomenex, Torrance, USA) (30m x 0.25mm i.d., d_f_ 1µm) in the GC was operated using the following conditions: split injection (4:1), initial oven temperature at 30 °C for 3.8 min, ramp at 30 °C min^-1^ to 100 °C. Helium was used as carrier gas at a flow of 0.7 mL min^-1^ constant flow.

High purity hydrogen gas (Alphagaz^TM^ 2 H_2_, Air Liquide, Düsseldorf, Germany) was used as the monitoring gas, also used to check the H_3_^+^ factor daily which was in the range of 2.39 to 2.43 ppm/nA during the measurement period. Two-point linear normalization of all *δ*^2^H values to the VSMOW scale were carried out using two working standards HUBG5 and HUBG3. The *δ*^2^H values of the working standards were calibrated against international reference substances (VSMOW2 [*δ*^2^H_VSMOW_ = 0.0 ± 0.3 mUr] and SLAP2 [*δ*^2^H_VSMOW_ = –427.5 ± 0.3 mUr]) by HTC-IRMS (IsoLab, Max Planck Institute for Biogeochemistry). The calibrated *δ*^2^H values in mUr vs. VSMOW were −191.7 mUr ± 0.8 mUr (n=9, 1σ) for HUBG5 and −272.9 mUr ± 1.5 mUr (n=11, 1σ) for HUBG3. For details of the calibration procedure, refer to the studies by Greule *et al*. (2019 & 2020).

Every tree sample was measured once followed by consecutive injections of both working standards after every sixth tree sample. Typical standard deviations of replicate quadruplicate analysis from one sample were in the range of 0.1 to 2.2 mUr (n = 4, 1σ). If necessary, an area drift correction was applied. Using Gaussian error propagation, the overall uncertainty in *δ*^2^H was in the range of 1.4 to 3.1 mUr.

### Definition of *δ* values and isotope fractionation

Stable hydrogen isotope results are reported relative to the VSMOW scale defined by the equation:

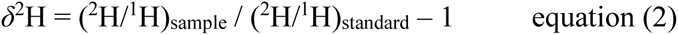

using the ‘delta’ (*δ*) notation. Following the suggestion by Brand & Coplen (2012) isotope *δ*-values are expressed in milli-Urey [mUr] (after Urey, 1948) instead of per mil [‰]. Hence, *δ*^2^H values that were formerly given as –250 ‰ for instance, are expressed as –250 mUr.

The apparent isotope fractionation between *δ*^2^H_TM_ and *δ*^2^H_precip_ (*ε*_TM/precip_) was calculated according to the following equation:

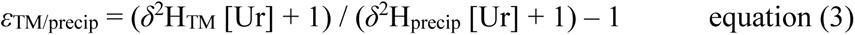

and is also reported in mUr.

### *δ*^2^H of precipitation and temperature data

The Online Isotope Precipitation Calculator (OIPC, accessible at http://www.waterisotopes.org/) using the IAEA database and interpolation algorithms developed by Bowen & Wilkinson (2002) and refined by Bowen & Revenaugh (2003) and Bowen *et al*. (2005), was used to calculate annual *δ*^2^H_precip_ values from wood sampling sites. Instrumental temperature data was received from the gridded time-series dataset CRU TS, version 4.07 at a resolution of 0.5° (Harris *et al*., 2020).

### Correction for elevational effects

In most regions of the world, the isotopic composition of precipitation is increasingly depleted towards heavier isotopes with rising altitude (Dansgaard, 1954). The study sites in this work cover elevations ranging from 45 to 3179 meters above sea level, necessitating the adjustment of *δ*^2^H_precip_ values (Fig S1). Initial *δ*^2^H_precip_ values were calculated using the OIPC model, which includes an altitude adjustment at a lapse rate of approximately –15.4 mUr per 1000 meters. However, a more recent study by Lehmann *et al*. (2022) employed a higher lapse rate of –22.4 mUr per 1000 meters to account for elevational effects. We applied a somewhat higher rate of –28 mUr per 1000 meters based on the following considerations:

The average adiabatic lapse rate is 6.5 °C per 1000 m (Kaimal & Finnigan, 1994; Stull & Ahrens, 1995) and (Greule *et al*., 2021) described a relationship between *δ*^2^H_precip_ and the mean annual temperature (MAT), by the following equation:

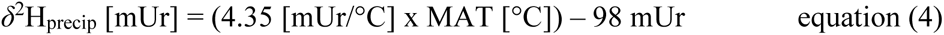

Using the adiabatic lapse rate and the relationship between MAT and *δ*^2^H_precip_ as shown in equation 4, a correction factor of –28 mUr per 1000 m of elevation was determined. Calculations of the apparent isotope fractionation between *δ*^2^H_TM_ and *δ*^2^H_precip_ values support the accuracy of the higher lapse rate applied in this study. While *ε*_TM/precip_ demonstrates a strong dependence on altitude concerning the uncorrected *δ*^2^H_precip_ values, it remains stable at a value of –185 mUr after correcting these values using a lapse rate of –28 mUr per 1000 m (Fig. S1). Although *ε*_TM/precip_ may vary with increasing depletion in *δ*^2^H_precip_ values, it is not expected to be affected by rising altitude.

To ensure consistency across the entire intercontinental scale, the European dataset was also corrected for the altitude effect. However, since the sample site altitudes in the European transect ranged only from 4 to 815 m asl, the altitude correction had minimal impact on *δ*^2^H_precip_ values from Europe, resulting in only a slight modification to the original data reported by (Greule *et al*., 2021) (Table S1).

### Statistics

Statistical analyses were conducted using R version 4.4.1 (R Core Team, 2024). Linear models were employed to assess the relationships between isotope values, apparent fractionation, latitude, and altitude, using the stats package in R. Significance levels were categorized as follows: highly significant (p < 0.001), significant (p < 0.05), and non-significant (p > 0.05). Tree core replicates per site along the American transect ranged from 4 to 16, while those along the European transect ranged from 2 to 10 (2021-1992). Please note that the number of replicates varies across the different time periods; further details can be found in table S1. For the comprehensive dataset evaluation, mean values from 13 and 25 locations were used for the American and European transects, respectively.

## Results and Discussion

### Spatial variations of *δ*^2^H_TM_ values of North American trees and relationship with precipitation

We measured 104 tree samples from 13 study sites across North America (Fig. 1) covering 90 years from 2021 to 1932. The tree cores were separated into three 30-years intervals representing the time periods 2021-1992, 1991-1962 and 1961-1932, respectively (for detailed overview see method section and Table S1). The most recent period (2021 to 1992) was further used to establish the relationship between *δ*^2^H_TM_ and modeled *δ*^2^H_precip_ values, which range from around –100 mUr to –200 mUr. We found a significant relationship between *δ*^2^H_TM_ and *δ*^2^H_precip_ values with a coefficient of determination of R^2^ = 0.61 (p = 0.0016) and a slope of the linear relationship of 0.52, indicating that the tree methoxy groups examined closely reflect precipitation isotope pattern at low to high latitudes of North America (Fig. 2, grey crosses, grey dashed line). Since the tree samples were collected from sites with highly variable altitudes (45 to 3179 m asl) we applied an altitude correction for *δ*^2^H_precip_ values received from the OIPC. Instead of using the standard correction factor of –15.4 mUr per 1000 meters typically applied by the OIPC, we implemented a factor of –28 mUr per 1000 meters of altitude. For a detailed explanation of the altitude correction, please refer to the Methods section (Correction for altitude effect). Using the altitude-corrected *δ*^2^H_precip_ values, R^2^ between *δ*^2^H_TM_ and *δ*^2^H_precip_ increases to 0.73 (p < 0.001) with a slope of 0.69 for the linear relationship (Fig. 2, red dots, red solid line).

**Figure 2.**
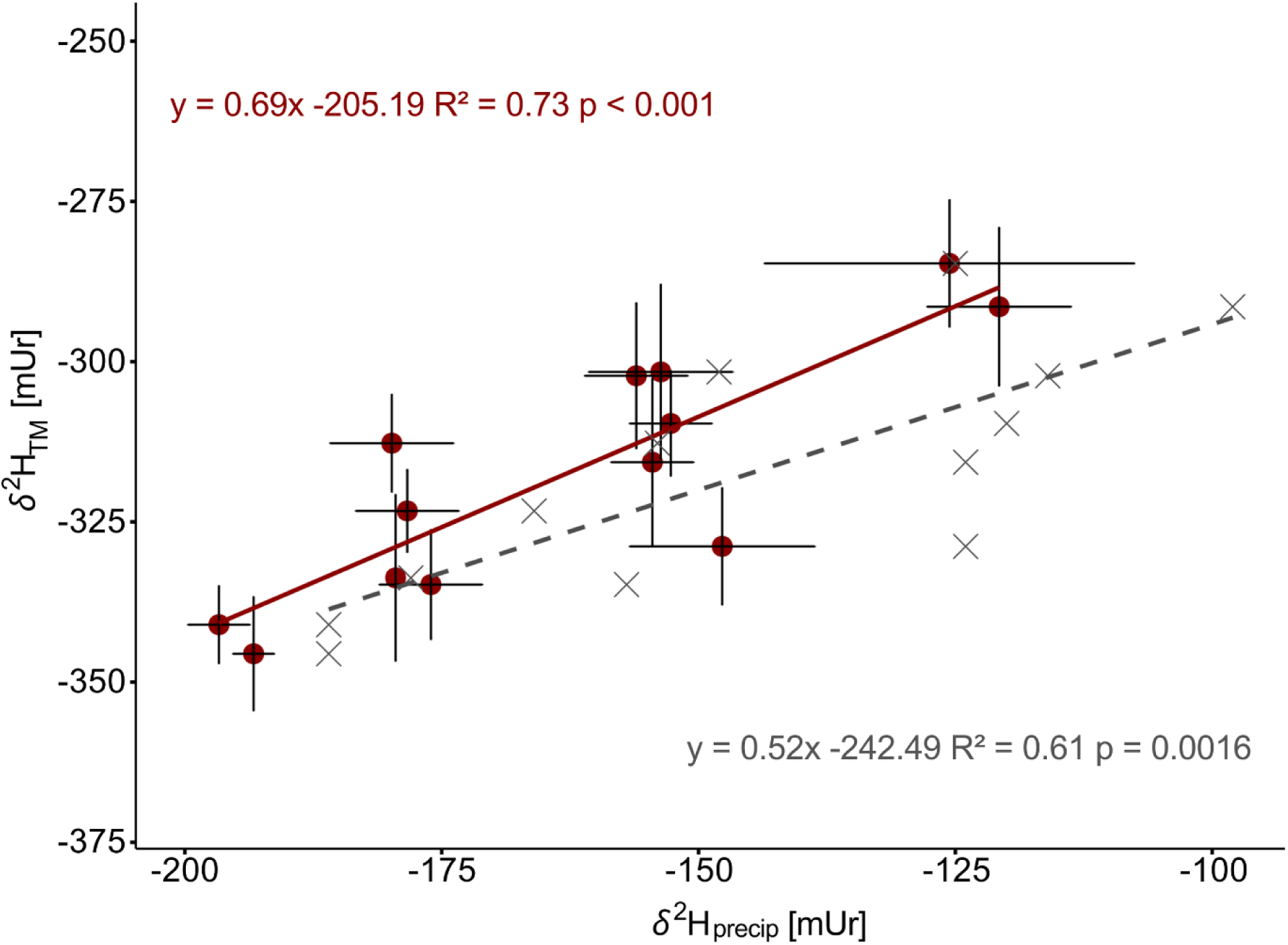
Average hydrogen isotope values (from 1992-2021) of tree methoxy groups at 13 sampling sites in North America and modeled precipitation (OPIC). Red dots represent elevation-corrected OIPC data and grey crosses the uncorrected *δ*^2^H_precip_ values, with red solid and grey dashed line showing linear regressions, respectively. The error bars on the y-axis represent the standard deviation (SD) among trees at each location (n ranging from 5-16), while the error bars on the x-axis indicate the 95% confidence interval of the modeled *δ*^2^H_precip_ values. The calculated mean apparent *ε*_TM/precip_ for the North American transect is –184.6 ± 12.4 mUr for the altitude-corrected data, and –201.5 ± 16.7 mUr for the uncorrected data (mean from all sample sites, see Table S1). These values are comparable to those from previous dataset from Europe, where *δ*^2^H_precip_ values ranged from –23 to –109, with corresponding mean *ε*_TM/precip_ of −200 ± 14 mUr and a linear relationship slope of 0.67 (R^2^ = 0.74) (Greule et al. 2021).

### Combing patterns of *δ*^2^H_TM_ values of North America and Europe - towards an intercontinental relationship of the Northern Hemisphere

In the following we have combined the results obtained in this study from North America with those previously reported for Europe (Greule et al. 2021) to establish an intercontinental relationship between *δ*^2^H_TM_ and *δ*^2^H_precip_ values across the Northern Hemisphere, covering temperate climates from lower to higher latitudes (36°N to 69°N). For both data sets modeled *δ*^2^H_precip_ values were corrected for altitude effects as mentioned in the section above and described in greater detail in the methods section. The combined data sets produced a strong and highly significant linear relationship (R^2^ = 0.94, p < 0.001) between *δ*^2^H_TM_ and *δ*^2^H_precip_ values with a slope of 0.71 ± 0.06 (Fig. 3). The calculated mean of *ε*_TM/precip_ of all tree wood samples from North America and Europe is −193 ± 13 mUr (individual sites ranging from −219 mUr to −162 mUr; see Table S1) and indicated by the dashed line in Figure 3.

**Figure 3.**
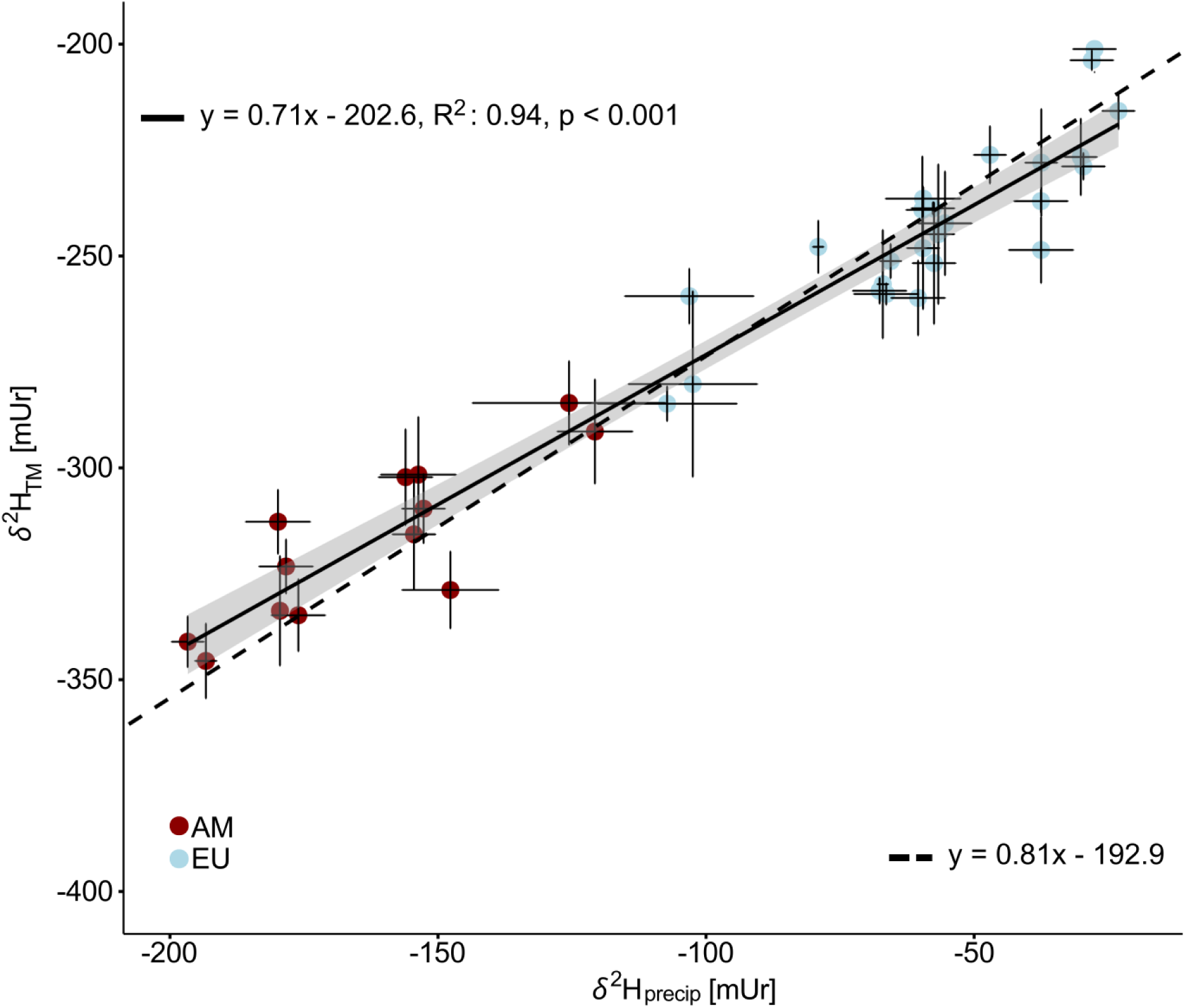
Hydrogen isotope values of recorded tree methoxy groups and modeled precipitation along North American (red) and European transects (blue). The solid line is a linear regression fit to the combined data and the grey shaded area the corresponding 95 % confidence band. The dashed black line indicates the mean constant fractionation of –193 mUr calculated from all tree sites in North America and Europe. Error bars on the y-axis represent the SD among trees at each location (n ranging from 2-16), whereas the error bars on the x-axis indicate the 95% confidence interval of the modeled *δ*^2^H_precip_ values. OIPC model results are corrected for elevational effects.

To explain the observed relationship between *δ*^2^H_TM_ and *δ*^2^H_precip_ values we draw on the findings discussed by Schmidt *et al*. (2003), Keppler *et al*. (2007) and Greule *et al*. (2021) regarding the systematics of ^2^H patterns in natural compounds, as well as the biosynthesis of plant methoxy groups and the corresponding stable hydrogen isotope distribution in their three hydrogen atoms.

The carbon precursor for the wood methoxy groups of phenylpropanoids (the monomeric units of lignin) originates from the amino acid serine. The central molecule involved in C_1_-metabolism is N^5^,N^10^-Methlyenetetrahydrofolate (Schmidt & Kexel, 1998). The methylene group (CH_2_-unit) originates from the C-3 position of serine and is transferred to tetrahydrofolate (THF) by the enzyme serine hydroxymethyltransferase. Studies have shown that the *δ*^2^H values of this CH_2_ unit are slightly depleted in ^2^H (up to 50 mUr) relative to the source water of plants (Zhang *et al*., 2002; Augusti *et al*., 2006). In the next step, the methylene group is reduced to a methyl group (CH_3_) by the NADH-dependent enzyme methylenetetrahydrofolate reductase (MTHFR), resulting in N^5^-Methyl-THF (Weilacher *et al*., 1996; Roje *et al*., 1999; Schmidt & Eisenreich, 2001; Schmidt *et al*., 2006). The reaction catalyzed by MTHFR also requires the non-covalently bound flavin adenine dinucleotide, which accepts reducing equivalents from NADH and transfers them to CH_2_-THF, forming the products NAD+ and CH_3_-THF (E. Trimmer, 2013). This transfer is associated with substantial ^2^H isotope fractionation (Schmidt *et al*., 2003) with depletions down to −780 mUr. (Billault *et al*., 2001; Schmidt *et al*., 2003; Martin *et al*., 2004) convincingly argue that the transfer of H⁺ from water to organic molecules typically requires photochemical reduction to H⁻, which is bound in the form of NADH. The theoretical isotopic fractionation associated with these processes can be estimated from the dissociation constants of H_2_O and D_2_O, yielding a hydrogen isotope fractionation between hydrogen gas and water (*ε*_H₂-H₂O_) of −645 mUr (Schmidt *et al*., 2003). In summary, the range of *ε*_H₂-H₂O_ values from biological H_2_ production spans from −550 mUr to −780 mUr. For further details on the biosynthesis of methoxy groups and hydrogen isotope fractionation, we refer readers to the study by Greule *et al*. (2021).

Based on the constraints related to the biosynthesis of tree methoxy groups, (Greule *et al*., 2021) proposed a simplified theoretical model for explaining *δ*^2^H_TM_ values. According to this model, the two hydrogen atoms in the CH_2_-unit are only slightly depleted in ^2^H relative to the source water (*δ*^2^H_precip_ values), with an apparent fractionation ranging from 0 to −50 mUr. In contrast, the third hydrogen atom, derived from NADH and incorporated into the methoxy group, shows a much larger apparent isotope fractionation relative to the source water, ranging from −550 to −780 mUr. However, the study by Greule *et al*. (2021) suggested that a fractionation of approximately −600 mUr for NADH is more likely than more depleted ^2^H values. Based on these assumptions, the *δ*^2^H_TM_ value for a specific *δ*^2^H_precip_ value can be calculated using the following equation:

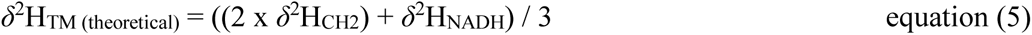

Using equation 5 with no fractionation between source water and the CH_2_ unit from serine, and a fractionation of −600 mUr for hydrogen derived from NADH, the theoretical *δ*^2^H_TM_ values can be estimated (*δ*^2^H_TM_modeled_) and compared with measured *δ*^2^H_TM_ values (Fig. 4). A strong linear relationship with an R^2^ = 0.94 (p < 0.001) and a slope close to 1 (1.06) supports the theoretical considerations proposed by Greule et al. (2021) to explain the empirical observations.

**Figure 4.**
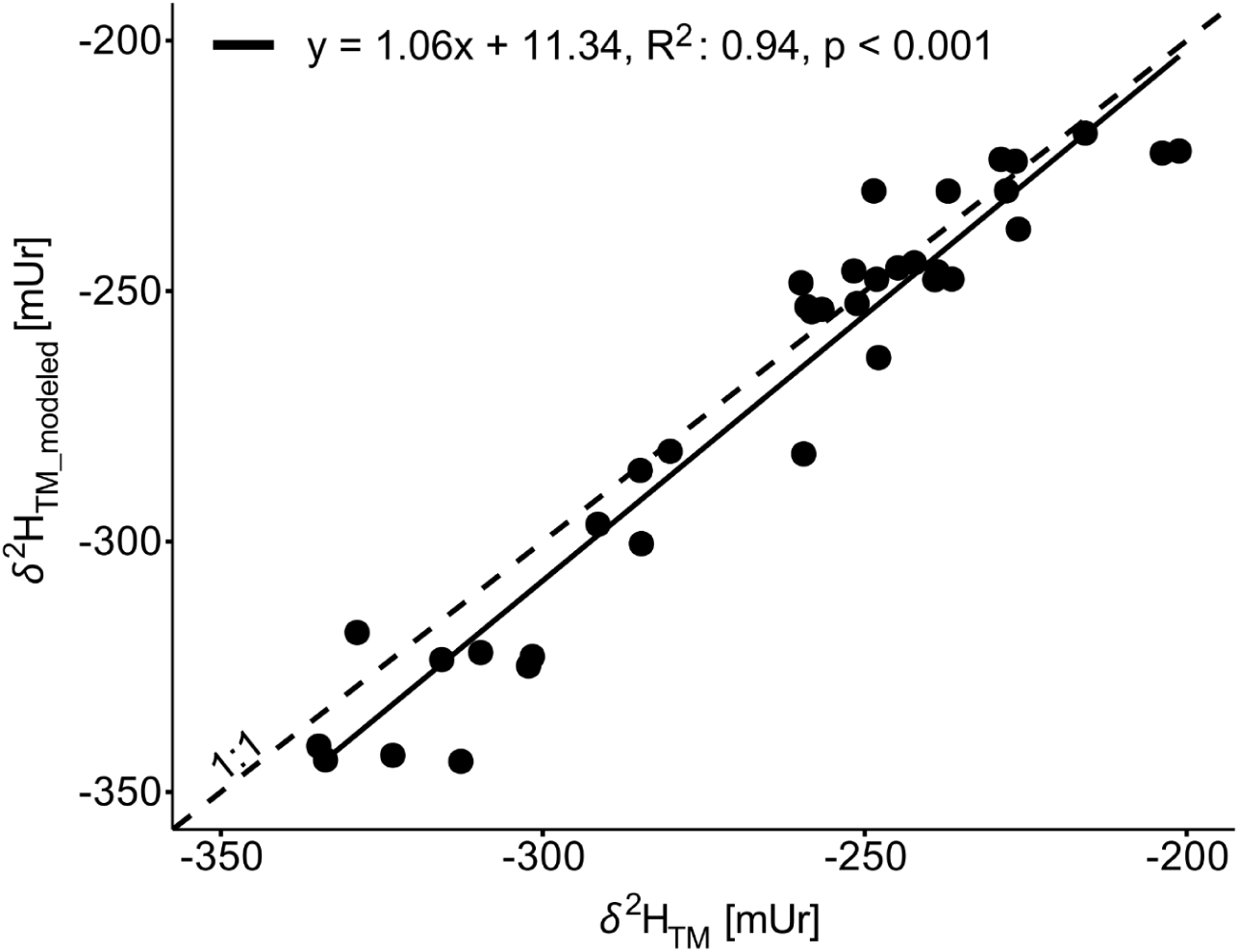
Modeled *δ*^2^H_TM_ values (*δ*^2^H_TM_modeled_) applying an apparent isotope fractionation between NADH and precipitation of –600 mUr and between the CH_2_ unit (C3 of serine) and precipitation of 0 mUr. Solid black line represents the linear relation between the modeled and the measured *δ*^2^H_TM_ values and the dashed black line shows the 1:1 relation.

Considering the theoretical constraints on *δ*^2^H_TM_ and *δ*^2^H_precip_ values, along with the strong alignment with empirical data (Figs. 3 and 4), equation (6) can be reliably applied to reconstruct *δ*^2^H_precip_ values, provided the *δ*^2^H_TM_ value of the tree sample is known

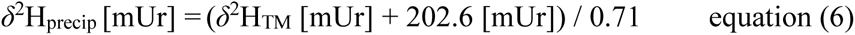

Alternatively, a uniform *ε*_TM/precip_ of –193 mUr (dashed line in Fig. 3) may be applied—which is commonly preferred in paleoclimate research—to calculate *δ*^2^H_precip_ for temperate regions of the Northern Hemisphere. However, additional factors at a single study site, such as soil water mixing, runoff, evaporation processes, or tree species specific varying water uptake zones, may also influence the isotopic signature of tree water sources (Lehmann *et al*., 2022). As a result, *δ*^2^H_precip_ may not be the sole or primary precursor of tree source water and, consequently, *δ*^2^H_TM_ values.

### Comparison of *δ*^2^H_TM_ values with wood cellulose and leaf n-alkanes

We further compare our results from the analysis of tree wood methoxy groups with other *δ*^2^H-based organic proxies related to trees, including wood cellulose and leaf n-alkanes, (Fig. 5). Recently, (Lehmann *et al*., 2022) compiled extensive tree cellulose data, encompassing measurements from diverse studies across North America, Europe, and Asia, covering various species and time periods (from the present day to 970 CE). This dataset was then compared with *δ*^2^H_precip_ data from the OIPC. Please note that these data were corrected for altitude effects by a lapse rate of −22.4 mUr per 1000 m (please refer to section methods “Correction for altitude”). N-alkane data were sourced from studies by Liu *et al*. (2019) and Liu *et al*. (2023) from various sites across China including Northwest China, the Tibetan Plateau, the Chinese Loess Plateau, Northeast China and South China. Site-specific *δ*^2^H_precip_ values were modeled based on data from 224 sites across China, provided by Wang *et al*. (2022). While tree cellulose exhibited minimal apparent fractionation relative to *δ*^2^H_precip_ values (−4.7 mUr) with a R^2^ of 0.69, n-alkanes showed substantial isotopic fractionation, in the range of −6 to −190 mUr and a weak correlation with *δ*^2^H_precip_ values (R^2^ = 0.11). While the isotopic signatures of n-alkanes and cellulose are affected by leaf water enrichment processes (Yakir & Sternberg, 2000; Sachse *et al*., 2010; Lehmann *et al*., 2022), *δ*^2^H_TM_ values are thought to directly reflect the tree xylem water which primarily represents water absorbed by the root system, which leads to the strongest correlations with *δ*^2^H_precip_ values (R^2^ = 0.94).

**Figure 5.**
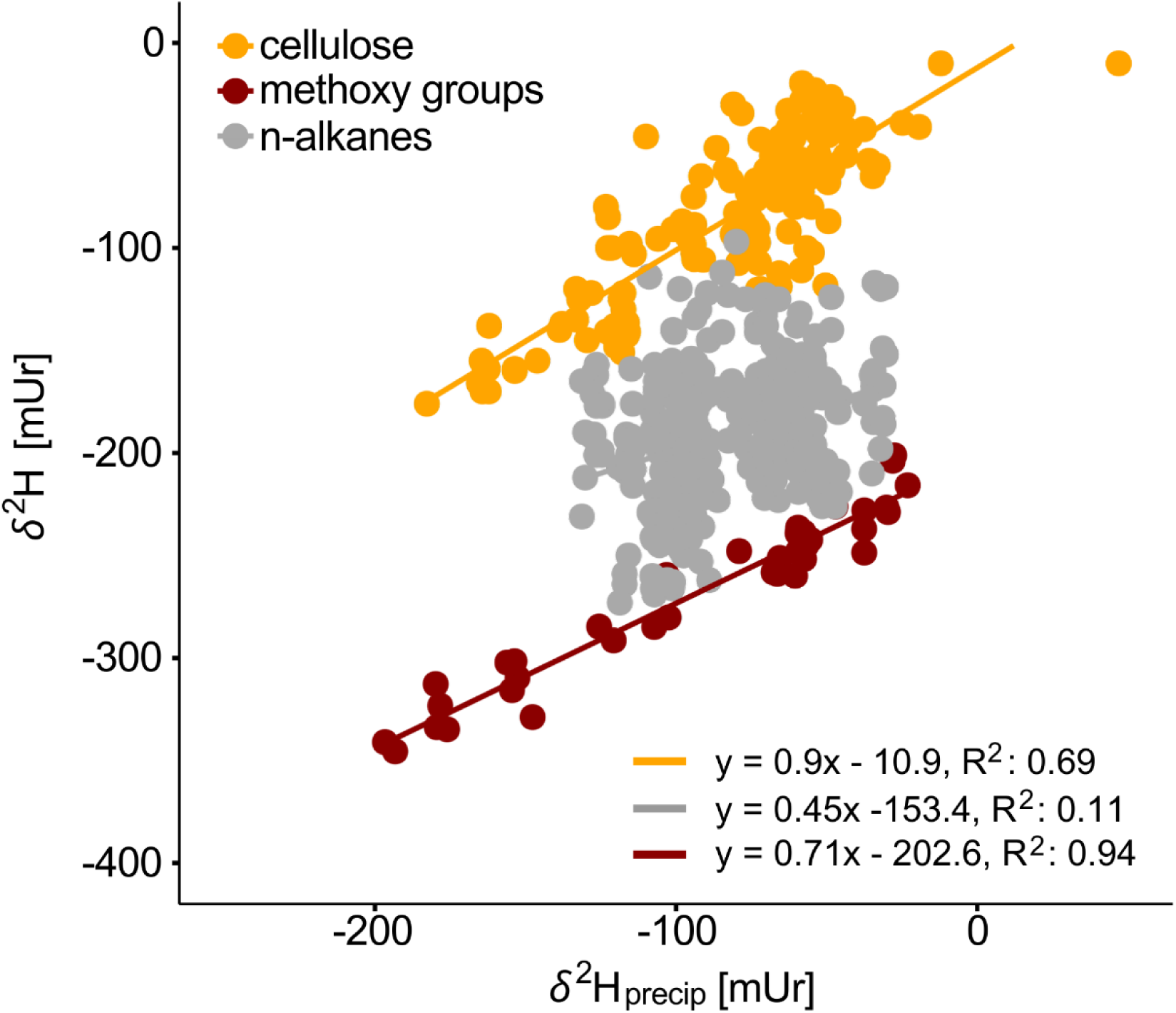
Comparison of *δ*^2^H values in tree stem methoxy groups (red) and cellulose (orange), together with n-alkanes from leaves (grey), plotted against modeled precipitation ratios for temperate regions of the Northern Hemisphere. Data for cellulose and n-alkanes were sourced from Lehmann *et al*. (2022) and Liu *et al*. (2023) and Liu *et al*. (2019), respectively

### Reconstruction of North American temperature changes since 1930

In temperate mid-latitudes, *δ*^2^H_precip_ values are predominantly influenced by air temperature (Dansgaard, 1964; Rozanski *et al*., 1993) and the robust relationship between *δ*^2^H_precip_ and *δ*^2^H_TM_ of tree ring samples can be applied to reconstruct temperature changes of mid-latitude regions on both spatial and temporal scales (Anhäuser *et al*., 2017; Greule *et al*., 2021; Porter *et al*., 2022; Wieland *et al*., 2022). Recent studies have applied temperature sensitivity rates of 2 to 4.5 mUr/°C to interpret changes in *δ*^2^H_TM_ and *δ*^2^H_precip_ values for temperature reconstructions over different periods. These applications include reconstructions from the past 100 years at Hohenpeißenberg, Germany; the Holocene and part of the Late Glacial period (up to 13,650 years BP) in the Eifel region, Germany (Anhäuser *et al*., 2014); and the Neogene (23-2.6 million years ago) in the Canadian Arctic region (Porter *et al*., 2022).

We analyzed the complete data set from North American trees to assess temporal changes in *δ*^2^H_precip_ values over a 90 year period (1932–2021), segmented into 30-year intervals (Table 1). We calculated the average temperature changes of each location and time interval relative to the most recent time period (2021-1992), providing an overall estimate for the North American continent. In the first step, we applied equation (6) to calculate mean *δ*^2^H_precip_ values from the measured *δ*^2^H_TM_ values for the most recent period (2021-1992). In the second step we, calculated mean *δ*^2^H_precip_ values from the measured *δ*^2^H_TM_ values for 1991-1962 and 1961-1932 intervals and determined the differences between these periods (Δ*δ*^2^H_precip_). Using the established relationship between *δ*^2^H_precip_ values and instrumental MAT (Harris *et al*., 2020) (equation 7 and Fig. S2) for the recent period 2021-1992, we applied a temperature sensitivity rate of 2.28 mUr/°C to convert Δ*δ*^2^H_precip_ into changes in mean annual temperature (ΔMAT) for the two prior periods, 1991-1962 and 1961-1932. These calculated changes were then compared with averaged instrumental temperature changes (Table 1).

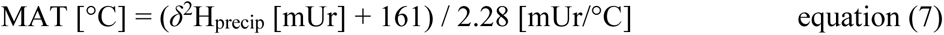

**Table 1:**
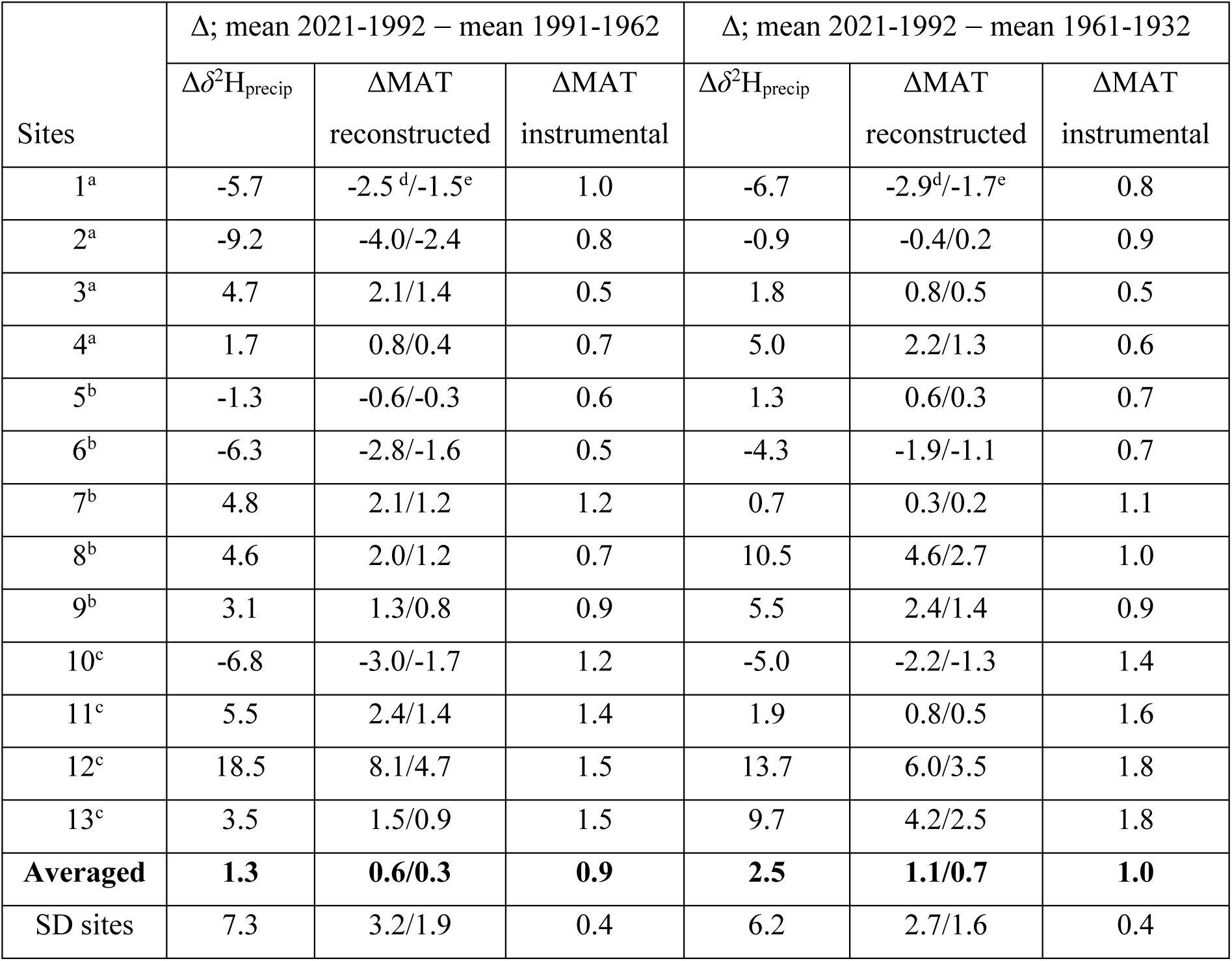
Reconstructed and observed temperature changes across 13 sites on North American continent (from 38° N to 69° N) relative to the reference period 2021-1992. ^a^sites 1-4, 38° N to 50° N; ^b^sites 5-9, 50° N to 65° N; ^c^sites 10-13, >65° N; ^d^temperature sensitivity rate of 2.28 mUr/°C, ^e^temperature sensitivity rate of 3.9 mUr/°C

For comparison, we also applied a higher temperature sensitivity rate of 3.9 mUr/°C, as recently used by Porter *et al*. (2022) to reconstruct Neogene temperature changes from wood samples in the Canadian Arctic Archipelago.

Reconstructed temperature changes across individual sites, relative to the periods 1991–1962 and 1961–1932, range from −4.0 to 8.1 °C and, in some cases, exceed those derived from instrumental records (CRU TS data). However, the average reconstructed temperature changes across all 13 North American sites are 0.6 ± 3.2°C and 1.1 ± 2.7 °C (for a temperature sensitivity rate of 2.28 mUr/°C) for the periods 1991–1962 and 1961–1932, respectively, closely aligning with observed data, which indicate changes of 0.9 ± 0.4°C and 1.0 ± 0.4°C.

When clustering North American sites into three latitude-based segments—sites 1–4 (< 50° N), sites 5–9 (50° N to 65° N), and sites 10–13 (> 65° N)—we found that the highest average temperature changes from 2021 to 1992, relative to 1961–1932, occurred at the highest latitudes (> 65° N). These increases were 2.2 ± 3.6 °C and 1.3 ± 2.1 °C for temperature sensitivity rates of 2.28 and 3.9 mUr/°C, respectively. At sites between 50° N and 65° N, these temperature changes were lower (1.2 ± 2.4 °C and 0.7 ± 1.4 °C, respectively). The smallest changes were observed at the lowest latitudes (< 50° N), with average changes of −0.08 ± 2.2 °C and −0.05 ± 1.3 °C. This pattern aligns with instrumental data, which show for the same time periods mean temperature changes of 1.65 ± 0.2 °C, 0.9 ± 0.2 °C, and 0.7 ± 0.2 °C for sites located at > 65° N, 50° N to 65° N, and < 50° N, respectively.

## Conclusions

Isotopic analysis of 104 North American trees from 13 sites between 38° and 69°N revealed a significant relationship between lignin *δ*^2^H_TM_ and *δ*^2^H_precip_ values. Given that *δ*^2^H_precip_ values in temperate mid-latitudes are primarily influenced by air temperature, this relationship was applied to reconstruct historical temperature variations. The *δ*^2^H_TM_-based temperature reconstructions for North America since 1932 align well with instrumental data, indicating that the greatest changes occurred at higher latitudes.

Comparing the North American evidence with data from European validates the recorded relationship between *δ*^2^H_TM_ and *δ*^2^H_precip_ values, demonstrating a strong and highly significant correlation (R^2^ = 0.94, p < 0.001) across a broad range of *δ*^2^H_precip_ values from −186 to −23 mUr) over a wide geographic transect (36.2° N to 68.8° N, 143,6° W to 24.2° E). This combined dataset, now encompassing 38 sites (13 in North America and 25 in Europe), supports a robust intercontinental estimate for temperate regions of the Northern Hemisphere revealing a fractionation between *δ*^2^H_TM_ and *δ*^2^H_precip_ values of –193 ± 13 mUr and supporting the reconstruction of historical *δ*^2^H_precip_ values from tree-ring *δ*^2^H_TM_ values. Furthermore, assessments of organic *δ*^2^H-based proxies from wood cellulose and leaf n-alkanes demonstrate that *δ*^2^H_TM_ values most accurately reflect variations in *δ*^2^H_precip_, primarily influenced by xylem water with minimal impact from leaf water. This finding presents unique opportunities for advancing climate and ecological research.

## CRediT authorship contribution statement

F.K and A.W. planned the study. J.E. and his team provided tree ring samples from North America, L.B. prepared the samples for isotope analyses and dendrochronological investigations and together with M.G. measured the stable hydrogen isotope ratios of tree methoxy groups. F.K., A.W. and L.B. drafted the manuscript with input from J.E. and M.G. All authors provided discussion and agreed to the final version of the manuscript.

## Data availability

All data supporting the findings of this study are available within this Article and its Supporting Information.

## Acknowledgement

We thank Philipp Römer, Inga Homfeld, Marcel Kunz, Eileen Kuhl, Davide Frigo and the Monostar team for sampling tree cores from North America. This research has been supported by the German Research Foundation (Grand nos. KE884/17-1 and ES161/12-1) and ERC Advanced Grant Monostar (AdG 882727).

## Supporting Information

**Fig. S1** Apparent isotope fractionation between hydrogen isotope values of tree methoxy group and precipitation.

**Fig. S2** Linear relationship between mean annual temperature and hydrogen isotope values of precipitation.

**Table S1** Sampling sites, isotopic and temperature data.

## Declaration of generative AI and AI-assisted technologies in the writing process

During the preparation of this work the author(s) used ChatGPT version 4.0 in order to rephrase text. After using this tool/service, the author(s) reviewed and edited the content as needed and take(s) full responsibility for the content of the publication.

