## Supplemental Figures for "Latitudinal gradients in stable hydrogen isotopes of North American trees"

The following Supporting Information is available for this article:

**Fig. S1** Apparent isotope fractionation between hydrogen isotope values of tree methoxy group and precipitation.

**Fig. S2** Linear relationship between mean annual temperature and hydrogen isotope values of precipitation.

**Table S1** Sampling sites, isotopic and temperature data (see Excel file)

**Fig. S1.** Apparent isotope fractionation between hydrogen isotope values of tree methoxy group ($\delta^2H_{TM}$) and precipitation ($\delta^2H_{precip}$) relative to altitude, shown before (grey crosses) and after (red dots) applying altitude correction to $\delta^2H_{precip}$ values. The black solid line and the grey dashed line represent the linear relationship.

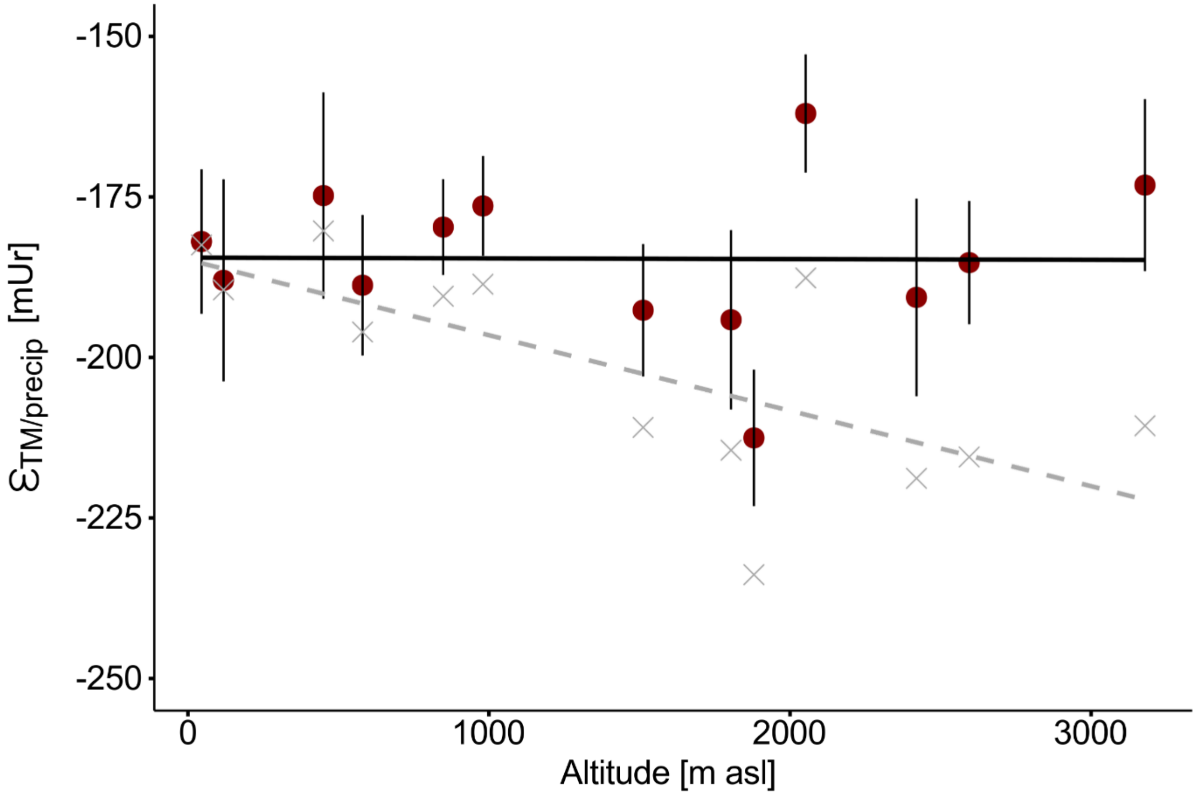

**Fig. S2.** Linear relationship between mean annual temperature (MAT, averaged from 1992 to 2021, received from the CRU TS network) and hydrogen isotope values of precipitation ($\delta^2H_{precip}$, received from the OIPC). Error bars on the y-axis indicate the 95% confidence interval of the modeled $\delta^2H_{precip}$ values, as calculated from the OIPC.

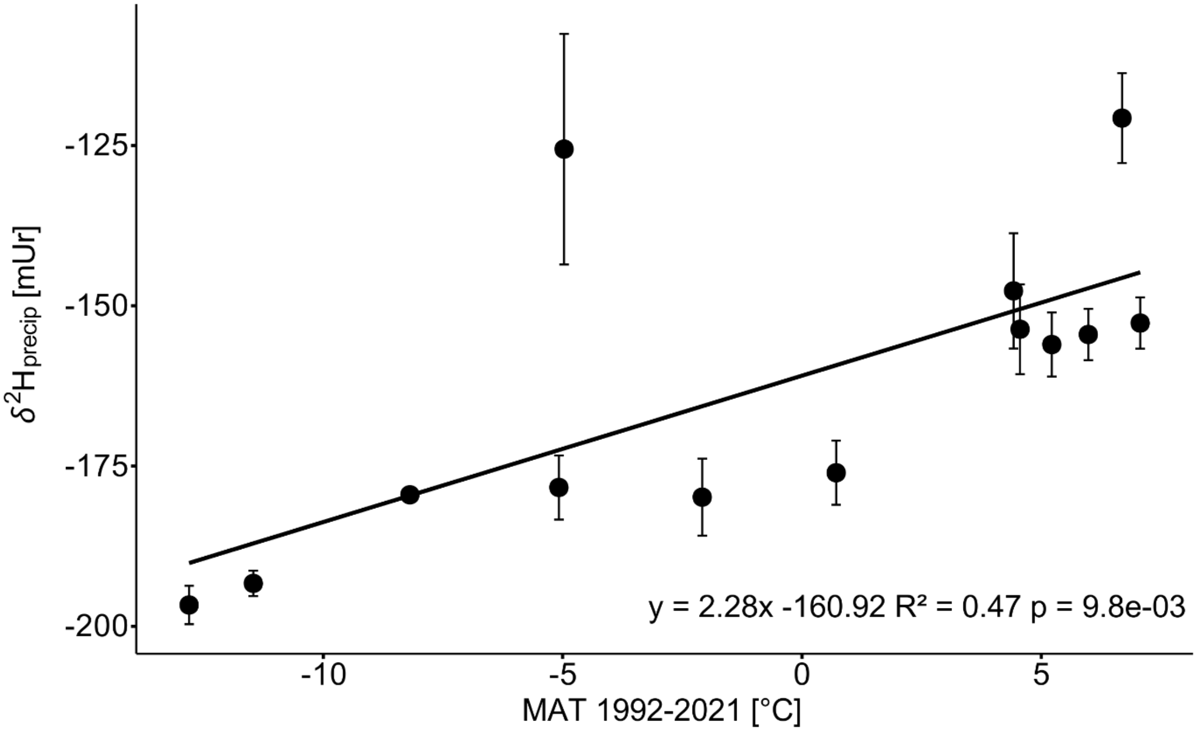
