## Supplemental Table 1 for "Latitudinal gradients in stable hydrogen isotopes of North American trees"

| | Site information | | | | 2021-1992 | | | 1991-1962 | | | 1961-1932 | | | | | | | Altitude corrected | | Temperature[a] | | |
|---|---|---|---|---|---|---|---|---|---|---|---|---|---|---|---|---|---|---|---|---|---|---|
| Continent | Site ID | Longitude | Latitude | Altitude | $\delta^2H_{TM}$ | SD $\delta^2H_{TM}$ | Replicates | $\delta^2H_{TM}$ | SD $\delta^2H_{TM}$ | Replicates | $\delta^2H_{TM}$ | SD $\delta^2H_{TM}$ | Replicates | $\delta^2H_{precip}$ | SD $\delta^2H_{precip}$ | $\varepsilon_{TM\_precip}$ | SD $\varepsilon_{TM\_precip}$ | $\delta^2H_{precip}$ | $\varepsilon_{TM\_precip}$ | 2021-1992 | 1991-1962 | 1961-1932 |
| America | 1 | -112,4 | 38,4 | 3179 | -302,2 | 11,3 | 4 | -298,2 | 13,0 | 5 | -297,4 | 11,8 | 5 | -116 | 5 | -211 | 13 | -156 | -173 | 5,2 | 4,2 | 4,4 |
| America | 2 | -120,3 | 39,4 | 2595 | -309,6 | 8,1 | 6 | -303,1 | 9,6 | 6 | -309,0 | 8,1 | 6 | -120 | 4 | -216 | 9 | -153 | -185 | 7,1 | 6,3 | 6,2 |
| America | 3 | -121,7 | 45,3 | 1804 | -291,4 | 12,3 | 6 | -294,8 | 10,8 | 6 | -292,7 | 12,1 | 6 | -98 | 7 | -214 | 14 | -121 | -194 | 6,7 | 6,1 | 6,1 |
| America | 4 | -114,3 | 45,9 | 2420 | -315,7 | 13,0 | 7 | -316,9 | 14,2 | 7 | -319,2 | 15,2 | 7 | -124 | 4 | -219 | 15 | -154 | -191 | 6,0 | 5,3 | 5,4 |
| America | 5 | -121,5 | 50,7 | 1880 | -328,8 | 9,1 | 10 | -327,9 | 9,1 | 10 | -329,7 | 10,1 | 8 | -124 | 9 | -234 | 10 | -148 | -213 | 4,4 | 3,8 | 3,7 |
| America | 6 | -117,2 | 52,2 | 2052 | -312,7 | 7,5 | 6 | -308,3 | 7,7 | 6 | -309,6 | 6,2 | 6 | -154 | 6 | -188 | 9 | -180 | -162 | -2,1 | -2,6 | -2,8 |
| America | 7 | -76,5 | 56,6 | 45 | -284,7 | 9,8 | 16 | -288,1 | 10,2 | 14 | -285,2 | 14,3 | 6 | -125 | 18 | -182 | 11 | -126 | -182 | -5,0 | -6,1 | -6,0 |
| America | 8 | -122,9 | 57,0 | 1512 | -334,8 | 8,5 | 9 | -338,1 | 8,7 | 7 | -342,3 | 10,7 | 5 | -157 | 5 | -211 | 10 | -176 | -193 | 0,7 | 0,0 | -0,2 |
| America | 9 | -134,5 | 58,4 | 450 | -301,6 | 13,6 | 12 | -303,8 | 11,7 | 11 | -305,5 | 10,6 | 11 | -148 | 7 | -180 | 16 | -154 | -175 | 4,6 | 3,7 | 3,7 |
| America | 10 | -145,9 | 65,4 | 980 | -323,3 | 6,4 | 5 | -318,5 | 15,8 | 6 | -319,7 | 13,7 | 6 | -166 | 5 | -189 | 8 | -178 | -176 | -5,1 | -6,3 | -6,5 |
| America | 11 | -133,4 | 68,3 | 118 | -333,7 | 12,9 | 8 | -337,6 | 15,1 | 7 | -335,1 | 12,1 | 7 | -178 | 0 | -189 | 16 | -179 | -188 | -8,2 | -9,6 | -9,8 |
| America | 12 | -141,0 | 68,7 | 580 | -345,6 | 8,8 | 9 | -358,7 | 10,5 | 8 | -355,3 | 10,7 | 7 | -186 | 2 | -196 | 11 | -193 | -189 | -11,5 | -13,0 | -13,2 |
| America | 13 | -143,6 | 68,8 | 848 | -341,0 | 6,0 | 6 | -343,5 | 5,6 | 6 | -347,9 | 4,1 | 3 | -186 | 3 | -190 | 7 | -197 | -180 | -12,8 | -14,3 | -14,6 |
| Europe | | -5,4 | 36,2 | 6 | -215,8 | 4,3 | 5 | | | | | | | -23 | 3 | -197 | 4 | -23 | -197 | | | |
| Europe | | -0,9 | 38,1 | 43 | -201,2 | 5,4 | 2 | | | | | | | -27 | 4 | -179 | 6 | -28 | -179 | | | |
| Europe | | -0,3 | 39,6 | 4 | -203,8 | 2,3 | 2 | | | | | | | -28 | 4 | -181 | 2 | -28 | -181 | | | |
| Europe | | -0,3 | 39,6 | 49 | -228,8 | 3,1 | 2 | | | | | | | -29 | 4 | -206 | 3 | -30 | -205 | | | |
| Europe | | 0,5 | 40,8 | 9 | -226,6 | 9,0 | 4 | | | | | | | -30 | 3 | -203 | 9 | -30 | -203 | | | |
| Europe | | -1,8 | 42,9 | 480 | -226,1 | 6,7 | 5 | | | | | | | -41 | 3 | -193 | 7 | -47 | -188 | | | |
| Europe | | 3,1 | 43,2 | 43 | -237,0 | 2,8 | 2 | | | | | | | -37 | 5 | -208 | 3 | -38 | -207 | | | |
| Europe | | 1,2 | 43,7 | 38 | -228,0 | 12,6 | 4 | | | | | | | -37 | 3 | -198 | 13 | -37 | -198 | | | |
| Europe | | 4,7 | 44,0 | 41 | -248,6 | 7,8 | 5 | | | | | | | -37 | 6 | -220 | 8 | -38 | -219 | | | |
| Europe | | 6,5 | 46,4 | 435 | -251,7 | 14,3 | 6 | | | | | | | -52 | 4 | -211 | 15 | -57 | -206 | | | |
| Europe | | 6,9 | 46,4 | 451 | -238,7 | 1,4 | 2 | | | | | | | -52 | 4 | -197 | 1 | -58 | -192 | | | |
| Europe | | 7,6 | 47,6 | 273 | -242,3 | 12,2 | 5 | | | | | | | -52 | 5 | -201 | 13 | -55 | -198 | | | |
| Europe | | 7,7 | 47,7 | 433 | -259,9 | 8,8 | 4 | | | | | | | -55 | 5 | -217 | 9 | -60 | -212 | | | |
| Europe | | 11,0 | 47,8 | 800 | -247,9 | 6,2 | 10 | | | | | | | -69 | 1 | -192 | 7 | -79 | -183 | | | |
| Europe | | 24,2 | 48,0 | 400 | -256,7 | 12,8 | 4 | | | | | | | -62 | 1 | -208 | 14 | -67 | -203 | | | |
| Europe | | 10,2 | 49,8 | 200 | -248,2 | 14,4 | 5 | | | | | | | -57 | 3 | -203 | 15 | -60 | -201 | | | |
| Europe | | 10,2 | 49,8 | 211 | -239,1 | 12,6 | 3 | | | | | | | -57 | 3 | -193 | 13 | -60 | -191 | | | |
| Europe | | 8,2 | 50,0 | 135 | -244,9 | 16,5 | 3 | | | | | | | -55 | 3 | -201 | 17 | -57 | -199 | | | |
| Europe | | 19,8 | 50,1 | 205 | -251,2 | 4,0 | 3 | | | | | | | -63 | 2 | -201 | 4 | -66 | -199 | | | |
| Europe | | 23,4 | 50,5 | 270 | -259,0 | 2,5 | 2 | | | | | | | -63 | 6 | -209 | 3 | -66 | -206 | | | |
| Europe | | 23,6 | 52,1 | 130 | -258,2 | 3,0 | 2 | | | | | | | -66 | 5 | -206 | 3 | -68 | -204 | | | |
| Europe | | -6,0 | 54,3 | 40 | -236,5 | 2,8 | 2 | | | | | | | -59 | 7 | -189 | 3 | -60 | -188 | | | |
| Europe | | 9,8 | 62,5 | 815 | -284,9 | 4,1 | 2 | | | | | | | -97 | 13 | -208 | 5 | -107 | -199 | | | |
| Europe | | 13,3 | 65,0 | 355 | -280,2 | 21,9 | 2 | | | | | | | -98 | 12 | -202 | 24 | -102 | -198 | | | |
| Europe | | 17,8 | 68,7 | 90 | -259,5 | 6,5 | 2 | | | | | | | -102 | 12 | -175 | 7 | -103 | -174 | | | |

[a] Mean annual temperature received from the CRU TS version 4.07 (Harris et al., 2020)
